# Multiscale Mechanisms of Human Memory Modulation by Deep Brain Stimulation

**DOI:** 10.64898/2026.04.14.718584

**Authors:** Yan Li, Ying Gao, Tong Li, Xiaojing Peng, Liang Zhang, Gangyao Yang, Liu He, Nikolai Axmacher, Tao Yu, Gui Xue

## Abstract

The highly variable effects of deep brain stimulation (DBS) on cognition highlight a fundamental gap in our understanding of how external perturbations interface with the brain’s ongoing, task-engaged state. The current study addressed this challenge by combining intracranial EEG recordings with hippocampal DBS stimulation in human participants during spatial sequence memory task. We found 50 Hz stimulation targeting hippocampal white matter enhanced memory performance, whereas 5 Hz stimulation targeting hippocampal gray matter impaired memory. These opposing behavioral outcomes were both governed by two dissociated mechanisms: a region-specific modulation of theta rhythms in an engagement-dependent fashion, and a global modulation of the fidelity of memory representations across the cortex. Our results suggest targeted stimulation can concurrently fine-tune specialized circuits and govern the brain’s global state, which not only offers a novel roadmap for understanding how external inputs shape human cognition, but also helps to design precise, state-dependent neuromodulation therapies.

## Introduction

Understanding how the brain’s activity causally constructs cognition—especially complex processes like memory—remains a defining challenge for neuroscience. To move beyond correlation and establish causality, a powerful strategy is to directly perturb the neural circuits, and observe how these interventions alter both neural dynamics and cognitive function. Deep brain stimulation (DBS) delivered via stereo-EEG enable us to simultaneously record and perturb neural activity in deep structures with millimeter precision, thus is increasingly used in cognitive neuroscience research^1^ to probe the causal underpinnings of cognition^2–4^, and the treatment of Parkinson’s disease^5–7^ and obsessive-compulsive disorder^8,9^. Despite their promise, the resulting effects have been strikingly variable. Memory, in particular, exemplifies this variability: stimulation has been shown to facilitate encoding in some studies^10–14^, but to impair or yield no measurable effect in others^15–20^. These discrepancies pose a central obstacle to clinical translation and underscore the need to clarify how exogenous perturbation interfaces with endogenous neural dynamics.

A primary determinant of variability lies in the stimulation parameters themselves. Frequency, amplitude, pulse width, and the anatomical locus of stimulation each exert profound influence on neural recruitment and downstream behavioral consequences^21–23^. The choice of frequency, for instance, can determine whether the stimulation serves to synchronize a target network or to functionally silence it^24–26^. Similarly, the precise anatomical placement of the electrodes determines which neural elements are primarily modulated, leading to fundamentally different network-level consequences^22,27,28^. Yet, most studies have examined these parameters in isolation, and little is known about their interactions. More fundamentally, the field still lacks an integrated causal framework tracing the pathway from stimulation input, through neural dynamics, to behavior^3,29^. Such a framework is essential if neuromodulation is to evolve from empirical trial-and-error to principled, mechanism-driven intervention.

Even more critical is the recognition that stimulation does not act on a passive circuit, but on one that is actively computing information. This leads to a crucial but often overlooked principle: the effects of stimulation are fundamentally *engagement-dependent*. For memory, its formation is not linked to a uniform brain state, but rather to a distributed and nuanced pattern of neural activity. For instances, successful memory encoding has been linked to both increases and decreases in neural activity across distinct brain regions, contingent on their specific functional roles^30,31^.This complexity suggests that for stimulation to be effective, it likely needs to go beyond simply targeting an anatomical location. A more successful approach may involve selectively engaging the specific neural populations functionally recruited by the task and modulating them in a manner that coordinates with their contribution to behavior. Although some work has examined connectivity-based predictors of stimulation efficacy^32–34^, the broader principles governing engagement-dependent and region-specific effects remain unknown.

A third, comparatively underexplored dimension concerns the impact of stimulation on neural representations—the population-level activity patterns that encode and differentiate mnemonic content^35–38^. Unlike localized oscillatory dynamics, representational fidelity reflects the coordinated engagement of distributed neural assemblies^39–41^. Whether and how DBS could alter these patterns may therefore represent a key undiscovered mechanism by which stimulation influences behavior.

While non-invasive studies have shown that stimulation can bias representational structure^42,43^, their limited spatial resolution constrains the ability to attribute these effects to specific anatomical regions. Intracranial EEG provides a rare opportunity to track representational changes across multiple spatial scales. It enables simultaneous measurement of stimulation-induced modulations in local oscillatory dynamics within defined circuits and in distributed neural representations spanning large-scale networks. This multiscale vantage point is essential for understanding how DBS reshapes the neural architecture that underlies human memory.

In summary, although both local oscillatory activity and distributed neural representations are known to support memory, it remains unclear whether they reflect distinct mechanistic pathways—and, critically, whether they causally mediate the behavioral effects of stimulation. In the present study, we utilized intracranial recordings in epilepsy patients to formally test this dual-pathway hypothesis. We systematically varied stimulation frequency and location during a spatiotemporal memory task, and employed multilevel mediation modeling to bridge neural and behavioral domains. This integrative approach not only addresses long-standing inconsistencies in the literature but also establishes a framework for advancing DBS from a powerful experimental tool toward a precise, mechanism-based therapeutic intervention for memory disorders.

## Results

### Stimulation effects on memory were modulated by site and frequency

Given the inconsistent behavioral effects induced by DBS^44,45^, our first objective was to identify stimulation protocols that reliably enhanced or impaired spatiotemporal memory. Participants performed a modified Corsi block-tapping test (Fig. 1a). In each block, two to three mini-sequences were learned against distinct color backgrounds. For each background, three locations were sequentially presented on a 4 × 4 grid and repeated three to six times during encoding while stimulation was delivered. Sequence length and repetitions were tailored to individual baseline performance. After a 10-s backward counting distractor, participants recalled the sequence associated with the cued colors. Stimulation targeted either the hippocampal gray matter or adjacent white matter tracts (Fig. 1b–c, Supplementary Table 1) at 5 Hz and 50 Hz (0.5 mA, 90 μs pulse width).

**Fig. 1.**
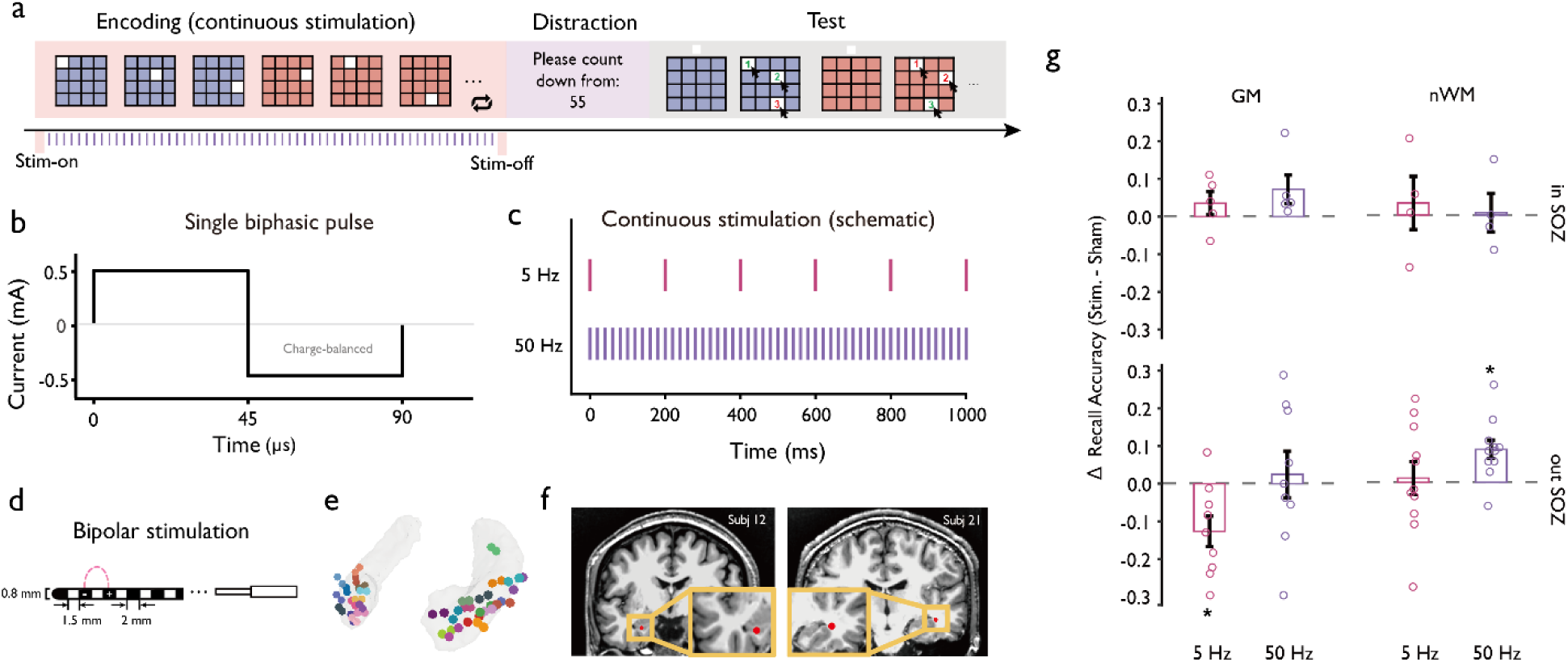
Experimental paradigm, stimulation protocol, and behavioral outcomes. **a,** Participants learned and recalled spatial sequences following a distraction interval. **b,** Schematic illustration of a single biphasic rectangular stimulation pulse with a duration of 90 μs and a peak amplitude of 0.5 mA. **c,** Schematic of the 1-s stimulation pulse sequence for 5 Hz and 50 Hz. **d,** Schematic diagram of depth electrode and bipolar stimulation configuration. Two adjacent contacts served as the anode and cathode, with current flowing between them. **e,** Hippocampal stimulation sites in MNI space, with examples of gray matter (**f**, left panel) and near white matter (**f**, right panel) sites (red dots = stimulation midpoints). **g,** Behavioral results. 5 Hz gray matter stimulation decreased memory performance, whereas 50 Hz near white matter stimulation increased the probability of memory recall. Each dot represents stimulation-induced change in recall accuracy for an individual participant. Error bars represent ±1 SEM to illustrate the variability of the group mean. Asterisks indicate conditions in which recall accuracy in the stimulation condition differed significantly from sham after false discovery rate correction. GM = gray matter; nWM = near white matter.

Based on prior work^14,44,46^, we hypothesized that gray matter stimulation would impair memory, whereas near white matter stimulation would enhance memory, and that these effects might be modulated by stimulation frequency. Consistent with our a priori expectation that stimulation-related modulation of memory would preferentially occur outside the seizure onset zone (SOZ), we initially focused behavioral analyses on stimulation delivered to regions outside the SOZ. A 2 × 2 analysis of variance (ANOVA) with stimulation site (gray matter vs. near white matter) and stimulation frequency (5 Hz vs. 50 Hz) revealed a significant main effect of site (*F*(1, 18) = 3.97, *p* = 0.0462, η² = 0.134) as well as a significant main effect of frequency (*F*(1, 18) = 10.15, *p* = 0.001, η² = 0.195). The interaction between site and frequency did not reach statistical significance (*F*(1, 18) = 1.162, *p* = 0.281, η² = 0.018).

Despite the absence of a significant interaction at the omnibus level, planned one-sample tests revealed a condition-specific pattern of stimulation effects. Stimulation at 50 Hz targeting near white matter significantly enhanced memory performance relative to baseline (mean difference = 0.0878; *t*(10) = 3.623; *p* = 0.005; FDR-corrected *p* = 0.0187; Cohen’s *d* = 1.092), whereas 5 Hz stimulation delivered to gray matter significantly impaired memory (mean difference = −0.126; *t*(8) = −3.126; *p* = 0.014; FDR-corrected *p* = 0.028; Cohen’s *d* = −1.042).

In contrast, no significant differences from baseline were observed for 5 Hz stimulation near white matter (mean difference = 0.011; *t*(10) = 0.262; *p* = 0.798; FDR-corrected *p* = 0.910; Cohen’s *d* = 0.079) or for 50 Hz stimulation delivered to gray matter (mean difference = −0.007; *t*(8) = 0.908; *p* = 0.909; FDR-corrected *p* = 0.910; Cohen’s *d* = −0.037). Besides, stimulation delivered within the SOZ did not show consistent or significant effects on memory across conditions (Fig. 1g), consistent with the hypothesis that pathological network activity may obscure stimulation-induced modulation.

Together, this pattern of results indicates that although the overall ANOVA did not reveal a significant interaction, stimulation effects on memory were highly condition-specific: memory enhancement was observed selectively during high-frequency stimulation near white matter, whereas memory impairment emerged selectively during low-frequency stimulation of gray matter.

### Theta power predicted spatiotemporal memory

To understand how stimulation modulated memory, we first had to establish the baseline neural signatures of successful spatiotemporal memory encoding. The role of theta oscillations in this process, known as the subsequent memory effect (SME)^47^, is notably nuanced. While numerous EEG/MEG studies have associated successful encoding with increases in theta power (theta SME+)^48–50^, a compelling body of intracranial evidence has linked successful encoding to theta power decreases (theta SME-)^47,51^. To replicate this, we compared theta power (3–8 Hz) between remembered and forgotten trials under sham condition to define the SME (Fig. 2a). Channels were classified as *theta SME+* (increased theta for remembered trials), *theta SME-* (decreased theta for remembered trials), or non-significant (non-SME) (Fig. 2a–b).

**Fig. 2.**
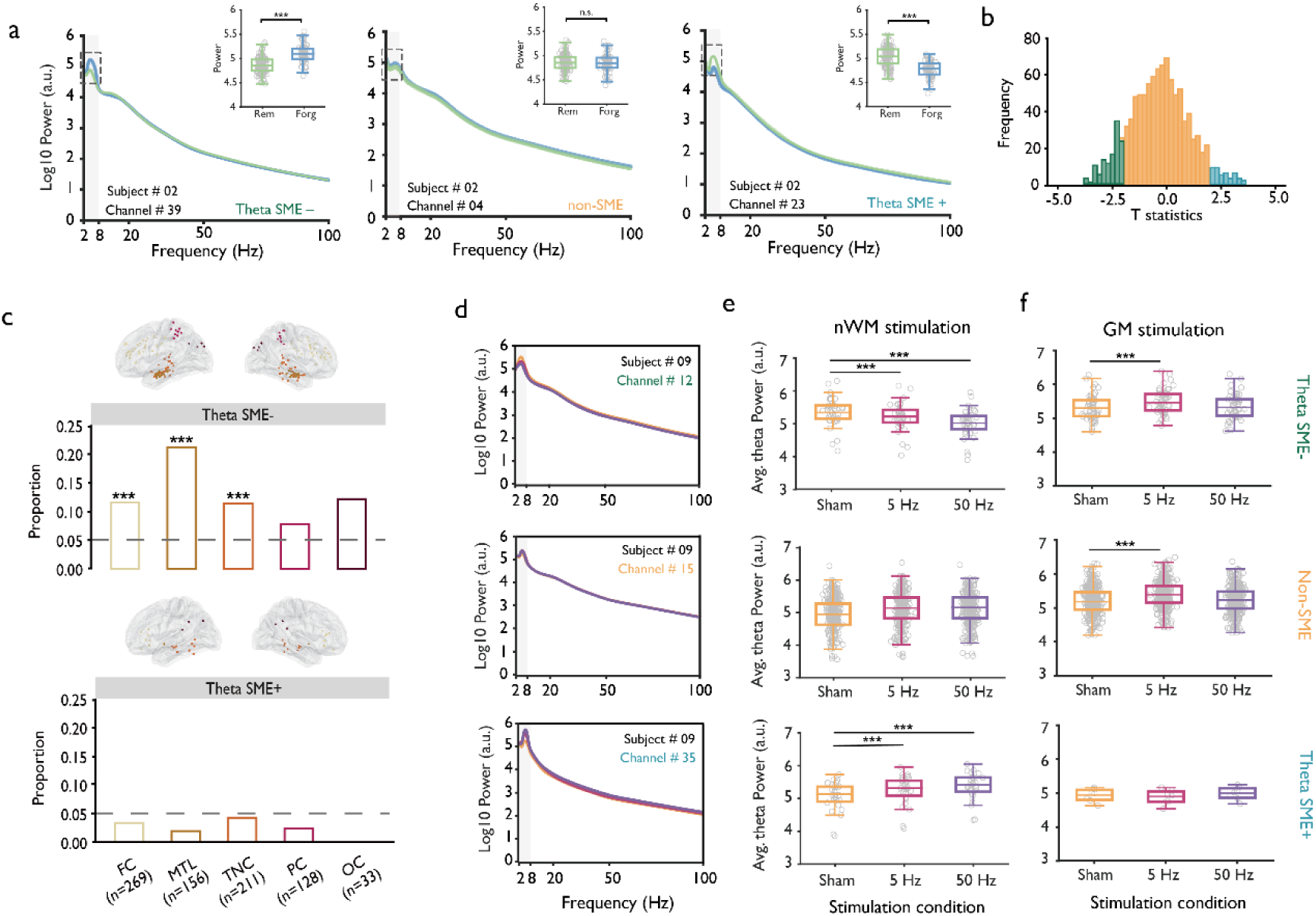
Stimulation-induced theta changes mirrored behavioral effects. **a,** Power spectral density was computed for each channel and each participant separately for correctly remembered (Rem.) and forgotten (Forg.) trials. Theta-band power (3–8 Hz) was averaged across trials within each condition, and trial-level variance was used in a paired t-test to identify channels showing a subsequent memory effect (SME). Channels with lower theta power for remembered trials were labeled as Theta SME- (e.g., Channel #39 in Subject #02, Left panel), whereas channels with higher theta power for remembered versus forgotten trials were labeled as Theta SME+ (e.g., Channel #23 in Subject #02, Right panel). The box plot in the top-left corner of the PSD shows the mean theta power (log10 scale) across trials for representative channels. **b**, Distribution of t values across channels. The histogram shows channel-wise t-values pooled across all subjects, with channels classified as theta SME- (green), non-SME (orange) and theta SME+ (blue). **c,** Proportion and spatial distribution of theta SME- (top panel) and theta SME+ (bottom panel) channels across ROIs. The gray dashed line indicates the expected false positive rate (p = .05). **d,** For a single subject receiving stimulation near white matter, the stimulation-induced power spectrograms are shown for three example channels: a Theta SME- channel (Top panel), a non-SME channel (middle panel) and a Theta SME+ channel (Bottom channel). **e, f,** Stimulation-induced theta power changes for different channel category under near white matter stimulation (**e**) and gray matter stimulation (**f**). Each dot represents an individual channel; the y-axis denotes the mean theta power (log10-scaled) for each channel. FC = frontal cortex; MTL = medial temporal lobe; TNC = temporal neocortex; PC = parietal cortex; OC = occipital cortex.

We found overall 12.80% channels (*n* = 102) were theta SME- channels, which was significantly higher than would be predicted by chance (binomial test, *p* < .001, Fig. 2b). By contrast, only 3.01% channels showed theta SME+ (*n* = 24, *p* = .998). The medial temporal lobe (MTL) contained the highest proportion of SME- channels (21.15%), especially in hippocampus (*n* = 21) and amygdala (*n* = 12). The number of theta SME-channels also exceeded chance level in frontal cortex (FC) and temporal neocortex (TNC) (binomial tests, both *p* < .001), but not in parietal cortex (PC) (*p* = .084) or occipital cortex (OC) (*p* = .055). The proportion of theta SME+ channels did not exceed chance in any ROI.

We also examined SME in the high gamma band, as some studies have reported high gamma power increases as a marker of successful memory encoding^10,52,53^. We found that 13.05% of channels (*n* = 104) were high gamma SME+ channels, which was above chance (*p* < .001, Supplementary Fig. 4a). This effect was observed across multiple ROIs (Supplementary Fig. 4c), suggesting that memory success was also associated with increases in high gamma power.

### Stimulation-induced theta changes paralleled memory outcomes

We next investigated whether the heterogeneous behavioral effects of stimulation on memory could be explained by changes in oscillatory power. If so, we would expect that the stimulation parameters that improved memory performance would further increase theta oscillatory power in theta SME+ channels and reduce theta power in theta SME-channels, and vice versa.

Supporting our hypothesis, we observed a significant three-way interaction among stimulation site, stimulation frequency, and channel SME status for theta oscillation (Χ*²* = 231.77, *p* < .001). In SME- channels, 50 Hz near white matter stimulation reduced theta (*t* = −16.26, *p* < .001, Fig. 2e, upper panel; also see Supplementary Fig. 3), while 5 Hz gray matter stimulation increased it (*t* = 18.69, *p* < .001, Fig. 2f, upper panel; also see Supplementary Fig. 3). In SME+ channels, 50 Hz near white matter stimulation increased theta (*t* = 11.58, *p* < .001, Fig. 2e, bottom panel), whereas 5 Hz gray matter stimulation had no effect (*t* = −1.76, *p* = .156, Fig. 2f, bottom panel). Modest effects were observed in non-SME channels as well. Besides, stimulation within the SOZ did not produce significant changes in theta power (Supplementary Fig. 5, all *p*s > .05), mirroring the inconsistent effect of stimulation in these areas on behavioral performance. High gamma and other frequency bands were unaffected by stimulation (Supplementary Figs. 4d, 6). Therefore, stimulation selectively modulated theta oscillations in ways that paralleled behavioral outcomes.

Previous work suggests that functional connectivity^32,34^, baseline excitability^54^, and cognitive states^55,56^ may also shape stimulation effects. To disentangle these contributions, we constructed a regression model that incorporated stimulation condition (5 Hz gray matter stimulation vs. 50 Hz near white matter stimulation), theta SME, functional connectivity, task-induced theta power, and distance to the stimulation site in order to quantify their respective contributions to stimulation-induced theta power changes at each recording site (Supplementary Fig. 7). These analyses showed that stimulation-induced theta changes were primarily determined by SME status, rather than connectivity, baseline activity, or anatomical proximity to the stimulation site.

### Theta power changes mediated stimulation effects on memory

The above analyses demonstrated that stimulation-induced theta power changes aligned with changes in memory performance, after controlling for multiple related factors. If theta modulation indeed constitutes the mechanism linking stimulation to behavior, a partial or full mediation effect would be expected.

We conducted a multilevel mediation model to test this hypothesis (see *Methods*). We found that 5 Hz gray matter stimulation impaired memory by increasing theta in SME-channels (indirect effect = −0.012, 95% CI = [−0.024, −0.004], *p* < .001; Fig. 3a).

**Fig. 3.**
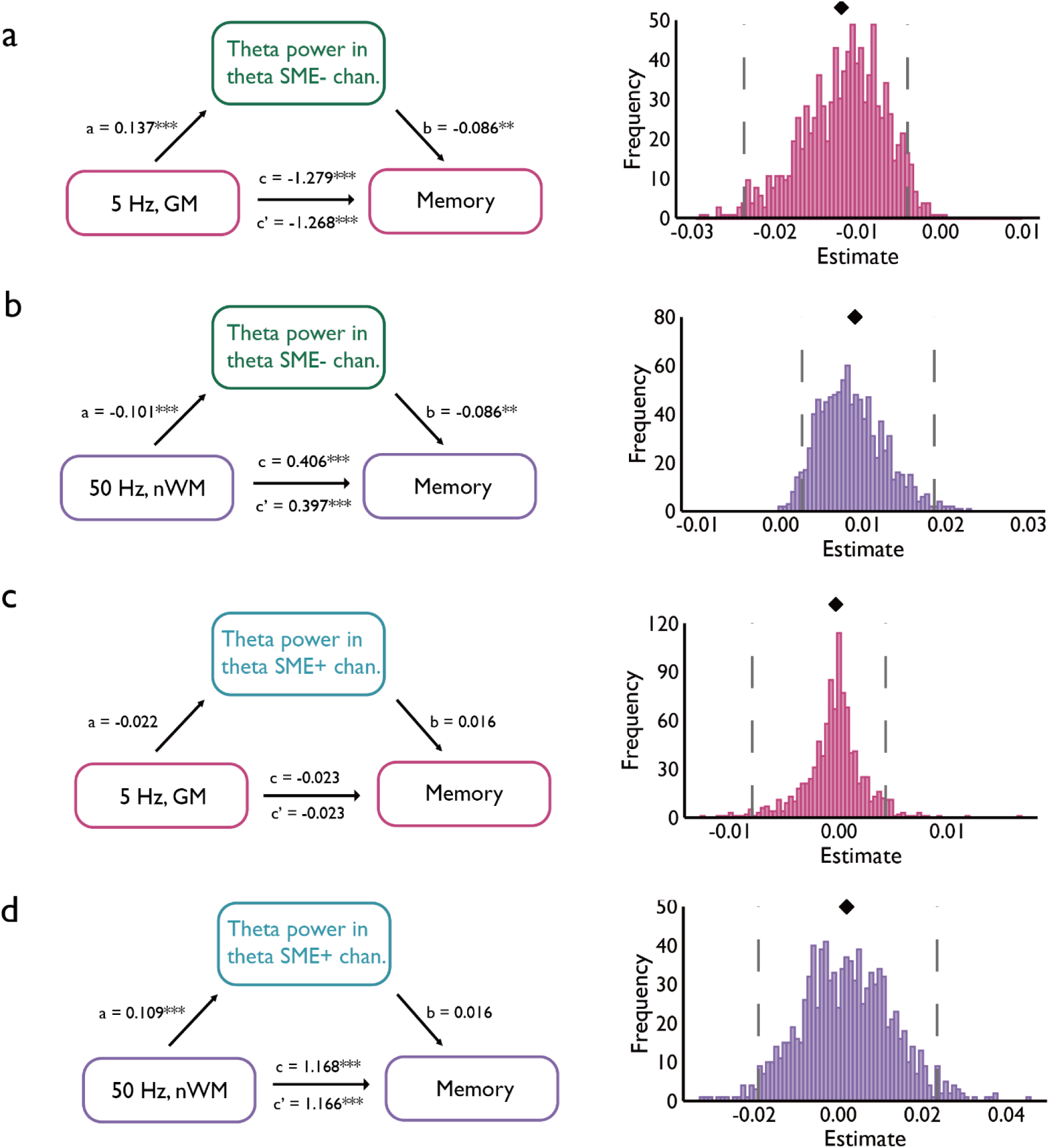
Mediation of DBS effects on memory by theta oscillations. Theta power in theta SME- channels mediated the effects of stimulation-induced memory performance under **(a)** 5 Hz gray matter stimulation and **(b)** 50 Hz near white matter stimulation. In contrast, no mediation effects were observed in theta SME+ channels under **(c)** 5 Hz gray matter stimulation and **(d)** 50 Hz near white matter stimulation. The histograms on the right show the bootstrap distribution (1,000 resamples) of the indirect effect (*a* × *b*). The black diamond above each histogram marks the observed indirect effect estimate, and the gray dashed lines indicate its 95% confidence interval.

Conversely, 50 Hz near white matter stimulation enhanced memory by reducing theta in SME- channels (indirect effect = 0.009, 95% CI = [0.003, 0.019], *p* < .001; Fig. 3b). No significant mediation was observed in SME+ channels (Fig. 3c–d). Multilevel mediation leverages the experimental design to identify theta activity as a mediator, providing mechanistic insight into how stimulation influences memory.

### Location-specific representations supported spatiotemporal memory

Our analyses thus far have revealed an oscillatory pathway, defined by task-relevant neural engagement as indexed by the SME^47^, that mediates behavioral outcomes. This mechanism, however, focuses on the neurobiological substrate of memory—the rhythmic activity of local circuits. Yet, a complete understanding of memory requires moving beyond the substrate to the level of information coding, i.e., the neural representations themselves. Memory is ultimately supported not just by rhythmic activity, but by the fidelity of distributed activation patterns that encode the specific content of an experience^36–38,40,57^. Prior studies using noninvasive stimulation have demonstrated enhancements in pattern similarity, suggesting that memory may be influenced by modulation of neural representations^42,43^. However, it remains unclear whether DBS of the hippocampus can influence memory by systematically altering these neural representations. To address this question, we investigated whether large-scale representational patterns could serve as neural signatures predictive of individual memory performance, and whether hippocampal stimulation induces systematic shifts in these global patterns during encoding.

We examined whether neural representations support spatiotemporal memory in the absence of stimulation. Spectral power from each channel was used to quantify representations (Fig. 4a), and three similarity measures were computed: within-location similarity (WLS), across trials sharing the same color background and location; within-context similarity (WCS), across trials with the same color background but different locations; and between-context similarity (BCS), across trials sharing neither color background nor location (Fig. 4b).

**Fig. 4.**
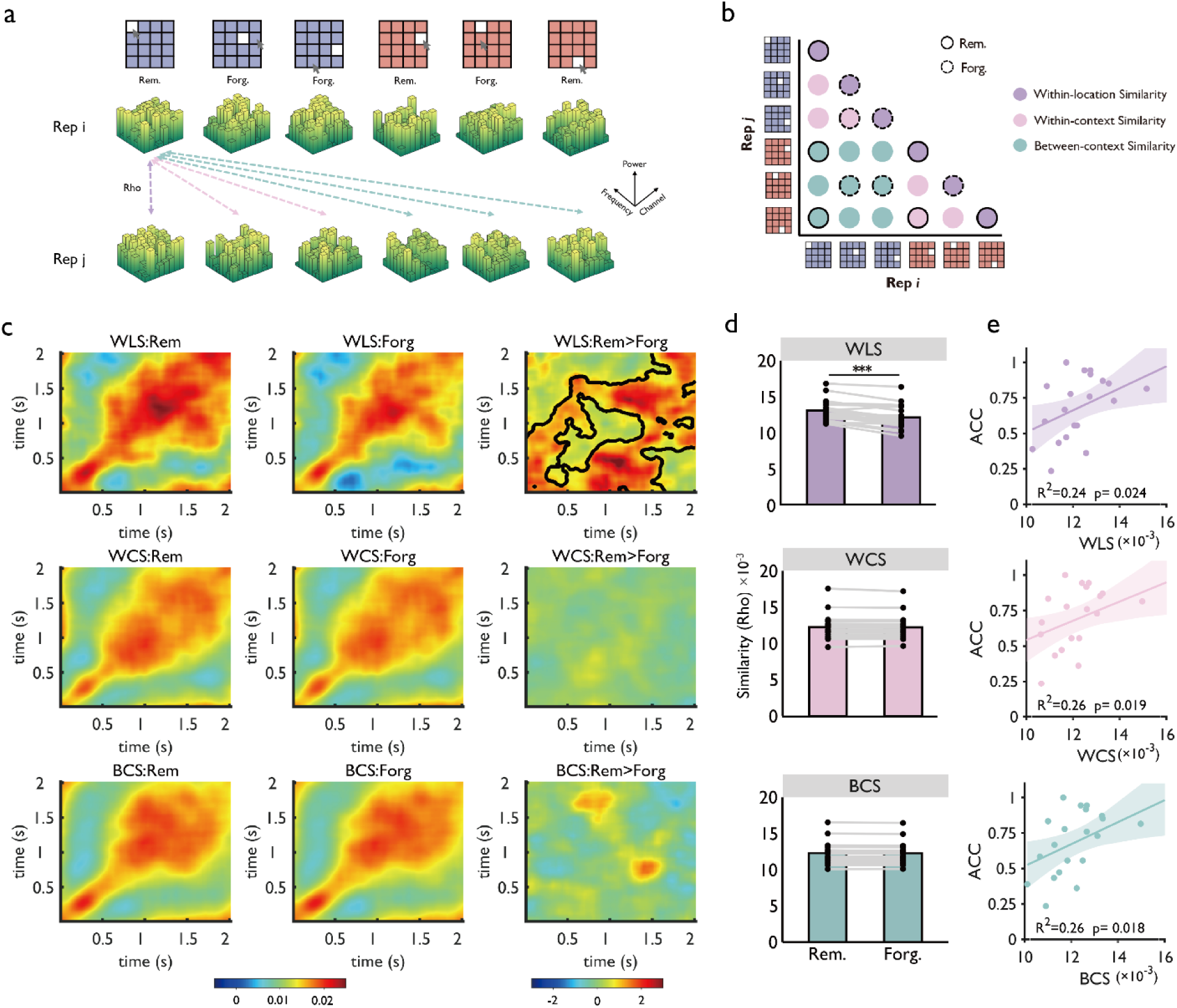
Within-location similarity (WLS) predicts spatiotemporal memory. **a,** Representational similarity analysis (RSA) was performed on time-frequency patterns from individual channels (2–100 Hz, 200-ms windows with 190-ms overlap). Correlations between trials were Fisher-z transformed. **b,** Schema of within-location similarity (WLS), within-context similarity (WCS) and between-context similarity (BCS). **c,** WLS (top), but not WCS (middle) or BCS (bottom), was greater for subsequently remembered than forgotten trials in the sham condition (cluster-based permutation, one-sided test). **d,** Across the entire encoding period, only WLS differentiated remembered from forgotten trials. Each dot represents data from one participant. **e,** WLS positively predicted individual memory accuracy after controlling for task difficulty; WCS and BCS showed no significant correlation.

Comparing subsequently remembered and forgotten trials revealed a significant SME for WLS (*t_sum* = 4503.3, one-sided *p* < .001, cluster-corrected; Fig. 4c, top panel), but not for WCS or BCS (Fig. 4c, middle and bottom panels). Across the encoding period, WLS again showed a significant SME (*t*(19) = 6.421, *p* < .001), whereas WCS (*t*(19) = 0.896, *p* = .381) and BCS (*t*(19) = 0.942, *p* = .357) did not (Fig. 4d).

Moreover, WLS positively predicted memory performance across participants (*t*(19) = 3.411, *p* = .003, Fig. 4e, top panel), after controlling for task difficulty (i.e., length and repetitions of mini-sequences), whereas WCS and BCS were not predictive (*p*s > .14). These results indicate that WLS constitutes a robust neural signature of spatiotemporal memory.

Jackknife analyses indicated that removing alpha- or beta-band features substantially reduced the WLS-related SME, whereas removal of theta-band features led to a comparatively smaller reduction. In contrast, removal of high-gamma features had minimal impact on SME (Supplementary Fig. 8). Therefore, this important finding suggests that theta and other frequency bands contribute to spatiotemporal memory via distinct mechanisms.

### WLS mediated the behavioral effects of stimulation

Having identified the neural representational indices for spatiotemporal memory, we next examined whether stimulation could modulate WLS, and thereby memory performance. Similar to the effects on theta oscillations, we found that stimulation modulated WLS in line with behavioral outcomes. Specifically, 5 Hz gray matter stimulation reduced WLS (*t* = −2.865, *p* = .014), whereas 50 Hz near white matter stimulation enhanced it (*t* = 3.413, *p* = .003; Fig. 5b). Mediation analyses revealed that WLS partially mediated the impairing effects of 5 Hz gray matter (Fig. 5c) and fully mediated the enhancing effects of 50 Hz near white matter stimulation (Fig. 5d). No mediation was found under other conditions (Fig. 5e–f). These findings highlight the critical role of WLS as a biomarker for memory and also as a mediator of stimulation effects.

**Fig. 5.**
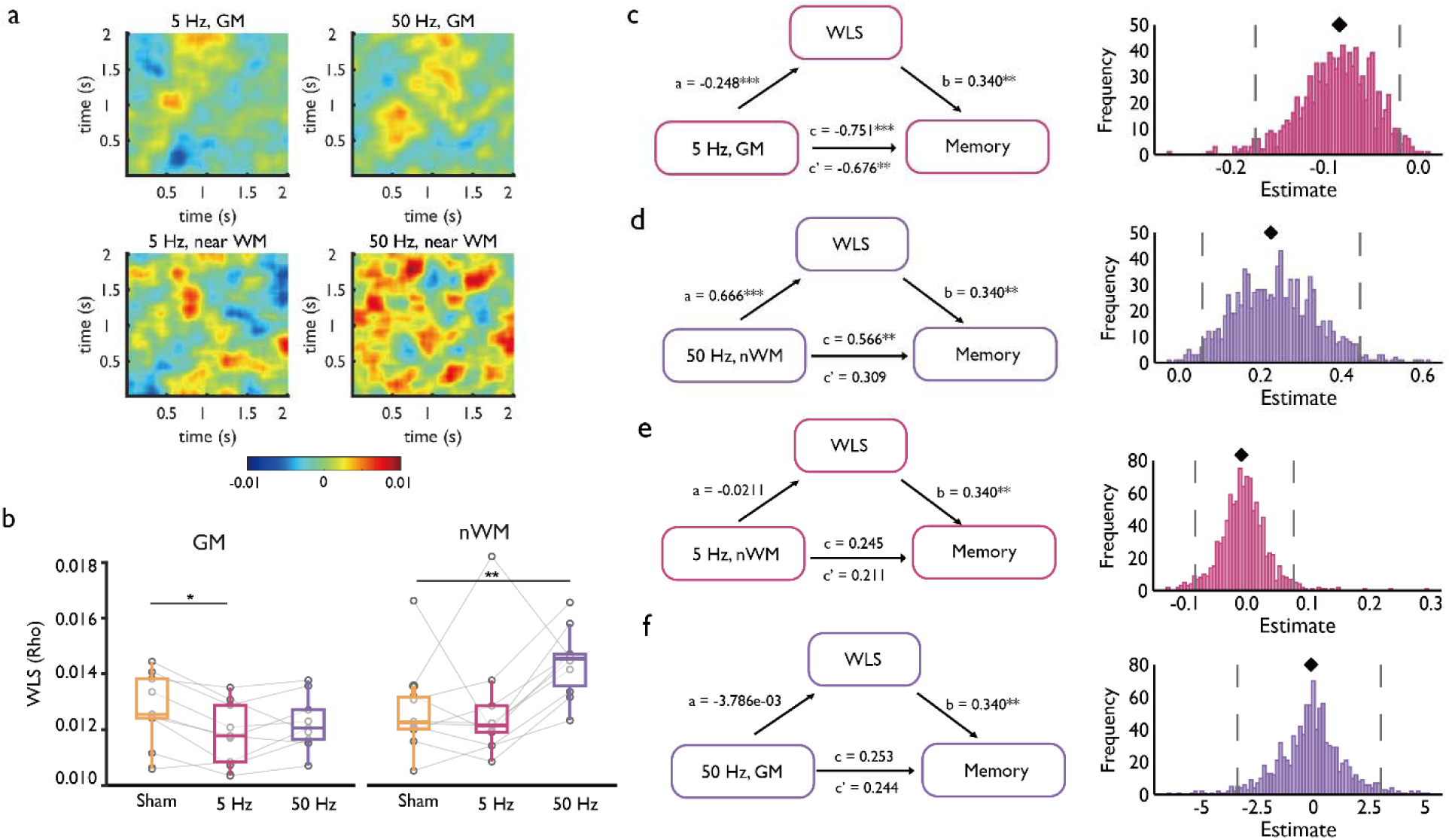
Location-specific neural representations mediate stimulation-induced memory changes. **a**, Stimulation-induced changes in WLS as compared to sham condition. WLS was computed using time-frequency features pooled across all channels within each location. **b**, Participant-level WLS for each stimulation condition. Each dot represents the mean WLS value obtained by averaging all entries of the within-location similarity matrix for a given participant under the corresponding condition. **c**, Mediation analysis showing that WLS partially mediates the negative effect of 5 Hz gray matter stimulation on memory performance. **d**, WLS fully mediates the positive effect of 50 Hz near white matter stimulation on memory enhancement. **e, f,** No significant mediation effects were observed for 5 Hz near white matter stimulation (e) or 50 Hz gray matter stimulation (f). Black diamonds mark the observed indirect effect estimate, and gray dashed lines indicate 95% confidence intervals.

### Anatomical localization of WLS_c_ SME

To localize the neural regions that contribute to representation-memory associations, we conducted two complementary analyses to quantify channel-level contributions to WLS SME. In the first analysis, we examined the WLS SME across trials for each channel separately for remembered and forgotten trials (channel-wise within-location similarity, i.e., WLS_c_; Fig. 6a, also see *Methods*). Channel-wise analyses revealed significant WLS_c_ SMEs in 10.66% of electrodes (*p* < .001), with significant proportions in PC, TNC, and MTL (Fig. 6b–c). In our second analysis, we employed a jackknife procedure to quantify how each individual channel contributes to the global WLS SME (see *Methods* and Supplementary Fig. 9a). Specifically, this method allowed us to assess the remaining global WLS SME after systematically removing the influence of each channel one at a time, thereby isolating its unique contribution. Consistent with the channel-wise analysis, we found that the proportion of WLS_c_ (jackknife) SME+ channels (8.41%) was significantly greater than the expected false-positive rate (binomial test, *p* < .001, Supplementary Fig. 9b). Again, these channels were mainly located in PC (*p* < .001), TNC (*p* = .037), and MTL (*p* < .001) (Supplementary Fig. 9d). Besides, there was a positive correlation across channels between the SME based on the single-channel WLS analysis and the channel-wise result from the jackknife analysis (*r* = .241, 95% CI = [0.097, 0.229], *p* < .001, Supplementary Fig. 9c) – showing that channels with prominent local WLS also contributed more to global WLS. Together, these results provide convergent evidence to suggest that neural representations in temporal and parietal regions support spatiotemporal memory.

**Fig. 6.**
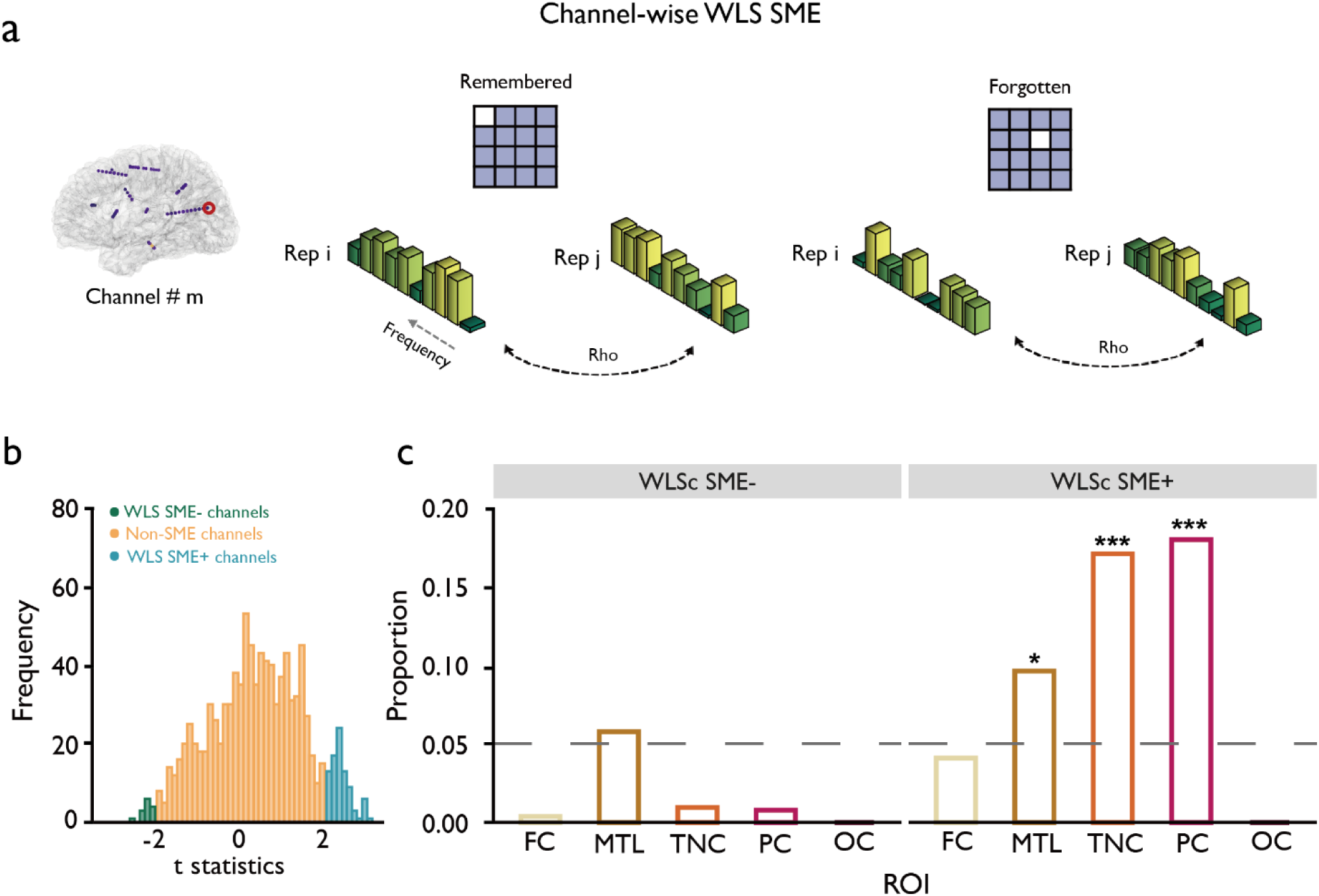
Channel-wise SME of within-location similarity. **a,** Schematic of channel-wise representational similarity analysis. For each channel, WLS_c_ (channel-wise within-location similarity) was computed under the sham condition between remembered and forgotten trials. **b,** Distribution of WLS_c_ SME t-statistics across channels. Channels were categorized as SME+ (significant positive SME), SME- (significant negative SME), or non-SME channels. **c,** Proportion of SME+ channels. The number of SME+ channels in PC, TNC, and MTL exceeded chance, whereas few SME- channels were observed.

### Global modulation of WLS by stimulation

We then examined whether stimulation-induced changes in WLS aligned with each channel’s contribution to WLS SME. Unlike theta oscillations, we found stimulation-induced WLS changes were more global and nonspecific (Fig. 7a). In particular, under 5 Hz gray matter stimulation, WLS was significantly reduced in both SME+ channels (*t* = −4.490, *p* < .001) and non-SME channels (*t* = −11.810, *p* < .001). In contrast, 50 Hz near white matter stimulation significantly increased WLS in both SME+ (*t* = 4.344, *p* < .001) and no-SME channels (*t* = 21.194, *p* < .001). Due to the very small number of WLS SME- channels, no reliable stimulation effects were observed for this category. Besides, stimulation-induced global changes in WLS were also observed when examining individual ROIs (Supplementary Fig. 10).

**Fig. 7.**
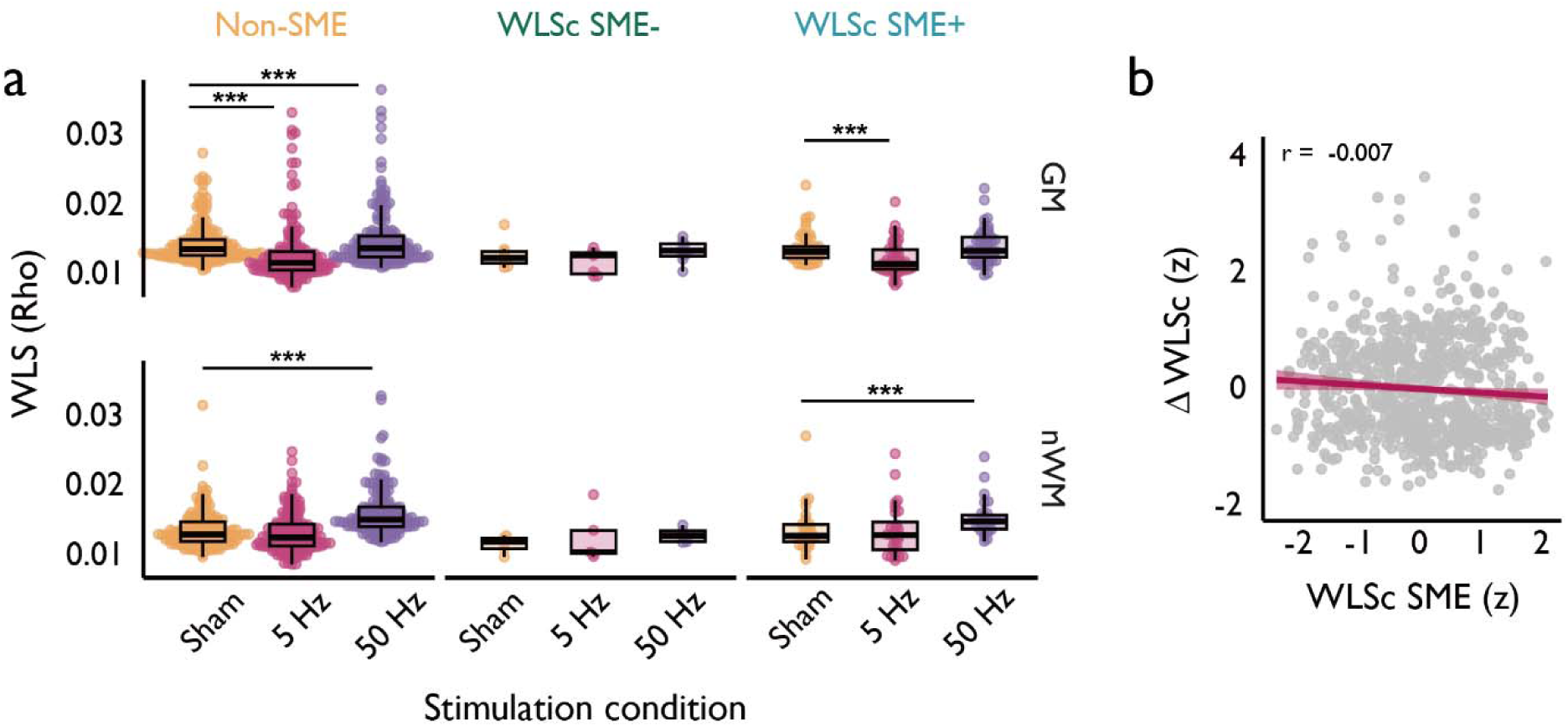
Global modulation of neural representations. **a,** Stimulation-induced WLS changes across channel categories. 5 Hz gray matter stimulation reduced WLS in both no-SME channels and SME+ channels, whereas 50 Hz near white matter stimulation enhanced WLS. **b,** Relationship between channel-wise WLS SME and stimulation-induced changes in WLS for each channel. Each dot represents individual channel. The x-axis shows the WLS_c_ SME t value for each channel, and the y-axis shows the stimulation-induced change in WLS, computed as WLS(stimulation) – WLS(sham). No significant correlation was observed, indicating that stimulation effects on WLS were not selectively stronger in channels that contributed more strongly to memory-related representations.

### Dissociated effects of stimulation on theta power and WLS

The above analyses together suggest that both theta oscillations and WLS mediated the stimulation effects on memory, yet their modulation patterns differed. Theta power changes were region-specific and depended on a given channel’s involvement in memory encoding, whereas WLS changes were global, affecting both SME-contributing and non-contributing channels.

To further characterize these dissociable effects, we directly compared the association between the modulating effect and task engagement. First, under 5 Hz gray matter stimulation, changes in theta power were negatively correlated with theta SME (*r* = −0.365, 95% CI = [−0.467, −0.263], *p* < .001). This indicates that the more a channel is functionally engaged in decreasing theta power for successful encoding, the more stimulation disrupted this pattern by increasing theta power. In contrast, changes in WLS were not related to WLS SME (*r* = −0.012, 95% CI = [-0.072, 0.047], *p* = .685). This suggests that the modulation of WLS was indiscriminate, occurring regardless of a channel’s specific contribution to memory. A formal bootstrap comparison confirmed this distinction: the predictive strength of theta SME on theta modulation was significantly greater than that of WLS SME on WLS modulation (mean difference = −0.352, 95% CI = [−0.460, −0.251], *p* < .001, Supplementary Fig. 11a).

Second, under 50 Hz near white matter stimulation, theta power changes were positively associated with channel SME (*r* = 0.201, 95% CI = [0.104, 0.298], *p* < .001), which implies that 50 Hz near white matter stimulation selectively amplified the endogenous memory-enhancing signals in functionally relevant channels. Conversely, WLS changes remained uncorrelated with WLS SME (*r* = 0.002, 95% CI = [-0.074, 0.078], *p* = .963), demonstrating a global, non-specific modulation pattern that was decoupled from a channel’s functional specialization. This dissociation was again confirmed by a significant difference in predictive strengths (mean difference = 0.199, 95% CI = [0.004, 0.431], *p* = .020, Supplementary Fig. 11b).

Finally, we assessed the relationship between the two memory-related signatures themselves. Across channels, the strength of a channel’s theta SME was not significantly correlated with the strength of its WLS SME (*r* = 0.063, 95% CI = [-0.015, 0.141], *p* = .115, Supplementary Fig. 11c). This statistical independence provides the evidence that these two neural signatures reflect genuinely distinct and parallel mechanisms.

Together, these results converge to demonstrate that hippocampal stimulation modulates memory via two independent pathways. It engages in a region-specific, engagement-dependent tuning of theta oscillations, while simultaneously enacts a global, non-specific regulation of distributed neural representations.

## Discussion

In this study, we provide a multi-level mechanistic account of how DBS modulates human spatiotemporal memory. Our findings reveal that the behavioral effects of DBS are co-determined by the specific parameters of stimulation and the brain’s intrinsic, functional state. We provide the first systematic behavioral evidence for an interaction between stimulation frequency and anatomical location. Critically, we go beyond describing the effects of DBS on memory performance and establish two distinct and dissociable mechanistic pathways that mediate these effects. First, we demonstrate a theta-specific oscillatory pathway, where stimulation’s impact is mediated by its modulation of theta power in a manner that is contingent on a channel’s functional engagement in the memory task. Second, we uncover a parallel representational pathway, where stimulation alters memory by globally modulating distributed neural patterns that encode information. Together, these findings contribute to an integrative framework that may help better understand prior inconsistencies in the field and inform the future development of more targeted, mechanism-driven therapeutic interventions.

Our results demonstrate that the behavioral effects of DBS depend critically on the frequency and anatomical placement of stimulation. Memory performance showed a statistically significant improvement when stimulation was delivered outside of the SOZ, proximal to white matter, and at higher frequencies. In contrast, low-frequency stimulation within intact hippocampal gray matter impaired memory encoding. This interference disrupts the fine-grained spike timing required for encoding^27^, consistent with reports that direct hippocampal stimulation is often deleterious for memory^18,46^.

Moving beyond *what* works, we investigated *how* stimulation affects memory. Our multilevel mediation analysis provides direct evidence that stimulation-induced memory changes are partially mediated by theta band oscillations. Prior work has linked theta to episodic memory encoding^47,51,58,59^ and shown that stimulation can modulate theta rhythms^24,26^. Our multilevel mediation analysis provides direct evidence that stimulation-induced memory changes are causally mediated by theta-band oscillations. The selectivity of this mechanism was striking: although high gamma power also exhibited a SME, its modulation neither tracked nor mediated behavioral outcomes, underscoring theta’s privileged role in memory. This does not diminish the importance of high gamma activity for memory formation itself, but rather underscores the specific role of theta oscillations as the primary mechanistic channel through which high-frequency stimulation influenced behavior.

This privileged role likely reflects the prominent role of theta oscillations for hippocampal computations. Theta rhythms provide a temporal scaffold for encoding and retrieval^47,60,61^, supporting phase coding that binds sequential inputs into coherent representations. The reason why stimulation preferentially modulates theta, in turn, likely stems from the intrinsic resonance of the hippocampal-entorhinal system, which is biophysically tuned to oscillate in the theta range^62^. Therefore, by acting on theta, stimulation is not just altering neural activity, but is intervening at the level of its core organizational structure.

This framework, which identifies theta modulation as the critical mechanistic link between stimulation and memory, is reinforced by observations within the SOZ. In these patients, the absence of a behavioral effect was accompanied by a corresponding neural silence: stimulation within the SOZ failed to elicit any significant modulation of theta power. This lack of neural responsiveness indicates that when the mechanistic link constituted by theta oscillations is compromised, the causal chain from stimulation to behavior collapses.

Another central contribution of our study is the establishment of an engagement-dependent principle for neuromodulation. We found that the behavioral and neurophysiological outcomes of stimulation were more strongly predicted by whether a channel exhibited an SME than by its anatomical location or even its resting-state connectivity. These findings support our conceptual framework of neuromodulation: the efficacy of DBS is not predetermined by parameters alone, but is contingent on the functional engagement of the targeted network in the cognitive process of interest. We empirically define this engagement using the SME, although we acknowledge that SME is not a pure measure of associative memory but may also reflect processes like attention, perception, and general task engagement that benefit later memory^47^. Thus, SME-defined channels may represent a holistic functional state where all necessary computations for successful encoding are aligned.

This engagement-dependent framework offers a unifying principle with broad implications. It likely explains state-dependent effects observed in other domains; for instance, in selective attention tasks, stimulation also preferentially modulates the attended (i.e., engaged) network^63,64^. This suggests that the most effective way to modulate a cognitive function is to engage the specific, distributed network that is actively implementing that function, providing a general law for understanding and optimizing a wide range of state-dependent neuromodulation phenomena.

A key conceptual advance of our study is the discovery of a second, parallel mechanism: the direct modulation of neural representations (WLS). This moves beyond oscillatory power to the level of information coding. Extant studies have established role of neural representational stability across repeated learning events in successful memory encoding^36,37,65^, and cognitive and neural manipulations^42,66,67^ can enhance pattern similarity and subsequent memory. Build on these studies, our study provides the first causal evidence that DBS can systematically enhance or impair spatiotemporal memory by acting on the stability of neural representational patterns. Specifically, low-frequency gray matter stimulation disrupted location-specific representations, whereas high-frequency white matter stimulation enhanced them, fully mediating behavioral outcomes. Intriguingly, a recent study has found that successful memory also benefits from lower neural similarity between distinct items, which reduces interference via pattern separation. Electrical stimulation can promote memory by facilitating temporal context drift and lowering inter-item similarity^68^. Together, these findings indicate that memory is supported by complementary neural representational mechanisms, which can be modulated by deep brain stimulation.

Unlike theta modulation, which was closely linked to channels showing an SME, the modulation of WLS was more global. The exact neural mechanisms underlying this stimulation effect require further examination, here we tentatively propose two possible mechanisms. First, whereas the propagation of neural oscillations is strongly constrained by network organization, the information transmission reflected in neural representations may be more diffusely distributed. Second, this global effect may originate from the influence of neuromodulators such as acetylcholine and norepinephrine, which broadly regulate cortical states reflecting attention, arousal, and signal-to-noise ratio^29,69,70^. Prior studies have demonstrated that fluctuations in attention and arousal exert strong effects on the stability of neural representations^71,72^, and future studies may investigate whether similar processes may account for the effects of DBS on memory (via neural representations). Taken together, these considerations suggest that WLS modulation reflects a more global state-dependent process, distinct from the network-specific modulation observed for theta activity.

We further demonstrate that neural representations and theta oscillations are mechanistically dissociated, pointing to distinct representational mechanism through which DBS can influence memory. Spectral analyses revealed that the WLS SME was driven by alpha, beta, and gamma activity, but not by theta band activity, underscoring a dissociation in terms of oscillatory frequencies. At the spatial level, the magnitude of theta SME did not correlate with WLS SME across channels, and stimulation-induced changes in theta power and WLS likewise exhibited uncorrelated spatial distributions. These dissociations indicate that oscillatory dynamics and representational stability reflect dissociable but potentially complementary aspects of memory encoding. Conceptually, our findings suggest that DBS can influence memory through at least two forms of neural modulation: one involving region-specific, engagement-dependent changes in theta oscillations that may scaffold temporal coding, and another involving more global changes in representational states that are associated with the stability of encoded content. These effects highlight the multifaceted nature of neuromodulation, in which local rhythmic processes and distributed representational dynamics may jointly contribute to cognitive outcomes.

Together, our findings reveal multiscale mechanisms through which DBS can simultaneously modulate network-level theta oscillations and influence global representational states. Conceptually, this underscores that neuromodulation is a context-sensitive process, contingent on both cognitive engagement and brain state—not merely a function of stimulation parameters or anatomical targeting. From a translational perspective, these insights support the design of more selective stimulation strategies aimed at engaging specific neural computations. This approach holds promise for developing targeted interventions—from supporting memory in epilepsy to modulating attention, decision-making, and learning in other neuropsychiatric conditions. Further research will be essential to verify the robustness of these mechanisms and to assess their therapeutic generalizability.

## Methods

### Participants

Twenty-five patients (mean age = 23.63 ± 8.20 years; 8 female) with medically refractory epilepsy undergoing surgical evaluation with stereoelectroencephalographic (SEEG) depth electrodes participated in this study. Detailed demographic and clinical information were provided in Supplementary information Table 1. The study protocol was approved by the Ethics Committee of Xuanwu Hospital, Capital Medical University, China (LYS[2020]074), and all participants provided written informed consent in accordance with the Declaration of Helsinki.

### Experimental design and stimulation protocols

Participants performed a modified Corsi block-tapping task designed to assess spatiotemporal memory (Fig. 1a). Under each stimulation condition, participants completed multiple sessions, with the number of sessions varying across participants due to clinical constraints (see Supplementary Table 2). Each session consisted of three blocks, and each block contained one sequence. Each sequence was composed of 2–3 mini-sequences, and each mini-sequence consisted of three spatial locations, leading to three trials per mini-sequence. Each sequence was repeated 3–6 times within a block to allow learning prior to recall. During encoding, three locations were sequentially displayed on a 4 × 4 grid (2s each). Following a 10s backward-counting distractor task to prevent rehearsal, participants recalled the sequence corresponding to a cued color.

During the recall phase, all grid locations were presented simultaneously, and participants reproduced the learned sequence by selecting locations sequentially using a mouse. To ensure consistent task engagement and avoid floor/ceiling effects, the number of mini-sequences and repetitions was titrated for each participant during a practice session.

During the whole encoding phase, continuous biphasic rectangular pulses (0.5 mA, 90 μs pulse width) were delivered at either 5 Hz or 50 Hz via a clinical stimulator (Fig. 1b–c, RISHENA, Jiangsu, China). During sham condition, no electrical current was delivered. To assess potential awareness of stimulation, participants were asked after stimulation sessions whether they experienced any sensation attributable to electrical stimulation.

All participants reported no subjective sensation, indicating effective blinding across stimulation and sham conditions. Stimulation targeted bipolar electrode pairs located either in hippocampal gray matter or adjacent white matter tracts (Fig. 1e–f). A trained clinician monitored physiological responses throughout all stimulation sessions to ensure patient safety. Additionally, a minimum of five minutes of seizure-free resting-state sEEG was recorded on a subsequent day to map intrinsic functional connectivity.

### Electrode localization

Stereo-EEG recordings were obtained using depth electrodes (Sinovation (Beijing); models: SDE-10, SDE-12, SDE-16). Each depth electrode was composed of 8 to 20 contacts, with an inter-contact spacing of 1.5 mm and a contact diameter of 0.8 mm (Fig. 1d). Following the procedure of channel localization described in^56^, each participant’s post-implantation CT image was first coregistered to their pre-implantation MRI, which was then spatially normalized to MNI space using Statistical Parametric Mapping (SPM12). The spatial coordinates of the channels were then rendered and inspected within 3D Slicer (https://www.slicer.org/). Pre-implantation structural MRIs were processed with FreeSurfer (surfer.nmr.mgh.harvard.edu) to generate cortical and subcortical parcellations. Contact locations were then assigned anatomical labels based on the Destrieux cortical atlas^73^. Adjacent channel pairs located within or in close proximity to the hippocampus were selected for bipolar electrical stimulation (Fig. 1f).

### Gray-white matter categorization

All stimulation sites were re-categorized as either located in the gray matter of the hippocampus or in nearby white matter, following previous studies^22,34^. Specifically, for each stimulation site, we first calculated the midpoint between the anode and cathode channels. A sphere with a radius of 4 mm centering on this midpoint was constructed. The gray and white matter vertices were identified based on segmentation performed in FreeSurfer, using each patient’s pre-implantation T1-weighted structural MRI. The median number of white matter vertices across all patients was 232 vertices. Participants were then grouped into gray matter and near white matter subgroups based on this median number.

### Data preprocessing and Time-frequency analysis

Intracranial EEG data was continuously recorded throughout the entire task periods using amplifiers from Brain Products (Brain Products GmbH, Germany) at a sampling rate of 2,500 Hz. All data preprocessing and analysis were performed using a combination of the FieldTrip toolbox^74^ and custom-written scripts in MATLAB (MathWorks, Natick, MA). We applied fourth-order Butterworth bandstop filter to remove 50 Hz line noise and its harmonic frequencies. Artifact rejection was performed using a two-step procedure. First, all channels were visually inspected in the time domain, and channels exhibiting obvious stimulation-related constant deflections were labeled.

Second, for each subject and each electrode, PSDs were computed and channels showing clear stimulation-related spectral contamination were removed (Supplementary Fig. 2). A total of 797 channels across 20 patients whose stimulation targets located outside the SOZ were retained for subsequent analyses. The average reference was applied to all clean channels.

The continuous neurophysiological data was segmented into epochs, starting from 3s before stimulus onset to 4s after the stimulus offset. Noisy trials contaminated by epileptic activities were also excluded after checking manually by Y. L. and an experienced clinician, Y. G. Time-frequency decomposition was performed using complex Morlet wavelets with six cycles across 40 log-spaced frequencies ranging from 2 to 100 Hz. The power of each frequency at each time point was log transformed. We only focused on the time period when the stimulus was presented.

For the resting state data, preprocessing steps were similar to the task iEEG data. After removing 50 Hz line noise, epileptic activities and artifacts were rejected through visual inspection. Averaged reference montage was applied before further analysis.

### Behavioral data analysis

For each participant, we calculated the mean recall difference between each stimulation condition and the sham condition (mean recall under stimulation − mean recall under sham). Analyses focused on out-SOZ participants to isolate the effects of stimulation outside the seizure onset zone. A 2 × 2 ANOVA was first conducted to assess the overall effect of stimulation site (GM and nWM) and stimulation frequency (5 Hz and 50 Hz) on memory recall. To determine whether stimulation reliably modulated memory, post-hoc one-sample t-tests were performed for each condition against zero. P-values from the post-hoc tests were corrected for multiple comparisons using the false discovery rate (FDR) method. This approach allows us to evaluate the effects of stimulation at the participant level while appropriately controlling for multiple comparisons.

### Identification of SME channels

Theta oscillations are critical for memory function, with several studies highlighting theta power as a potential neural marker for SME^51,75^. To identify channels associated with successful memory encoding, we focused on spectral power changes within the 3–8 Hz frequency band, which has been implicated in memory-related processes. For each channel, we first averaged the 3–8 Hz band power separately across remembered and forgotten trials. A trial was considered as remembered only when the participant selected the location and its position in the sequence correctly. A trial was considered forgotten if either the location or its temporal order was incorrect. T-test was then conducted at channel level in the absence of stimulation (Fig. 2a).

We categorized the channels into three groups based on the direction and significance of the observed SME: *theta SME+ channels*, *theta SME- channel*s and *non-SME channels*. *Theta SME+ channels* referred to channels that exhibited significantly greater theta power during successfully remembered trials compared to forgotten trials. *Theta SME-channels* were those that showed significantly lower theta power during remembered trials compared to forgotten trials. *Non-SME channels* were defined as channels that did not show significant SME.

To quantify stimulation-induced changes in theta activity, theta power was computed at the channel level for each stimulation condition separately. For each channel, 3–8 Hz band power was estimated during the encoding period under each stimulation condition (5 Hz stimulation, 50 Hz stimulation, and sham), using the same spectral analysis pipeline as described above. Statistical comparisons of theta power were performed across stimulation conditions at the channel level, with channels grouped according to their SME classification (theta SME+, theta SME−, and non-SME).

### Region of interest (ROI) analysis

In order to determine whether the SME channels were preferentially localized to specific brain regions, we performed an ROI analysis. ROIs were defined based on the anatomical locations of the implanted channels (Supplementary Fig. 1). Specifically, we selected five major ROIs: frontal cortex (FC, *n* = 269, e.g., primary motor area, premotor area, supplementary motor area, prefrontal area), medial temporal lobe (MTL, *n* = 156, hippocampus, amygdala, parahippocampal gyrus), temporal neocortex (TNC, *n* = 211, e.g., superior temporal area, middle temporal area, inferior temporal area, temporal pole), parietal cortex (PC, *n* = 128, e.g., postcentral area, inferior parietal area, superior parietal area, supramarginal area, angular area, lingual area, precunes), and occipital cortex (OC, *n* = 33, e.g., inferior occipital area, middle occipital area, superior occipital area, cuneus). For each ROI, we quantified the proportion of SME channels and tested whether this exceeded chance levels using a binomial test.

### Representational Similarity Analysis

Representation similarity was estimated between trials across repetitions (Fig. 4a). We used 200-ms sliding windows, with a step of 10ms, to compute the representational similarity. Within each time window, spectral power was averaged across all time points for each frequency and each channel, producing a feature vector representing the neural activity pattern across both frequencies and electrodes. The individual’s representation similarity was identified by computing the spearman’s correlation between two trials with vectorized spectral power features, and the correlation value was Fisher Z-transformed^57^. Based on trial conditions, we further categorized representational similarity into three types: *within location similarity (WLS)* was calculated between trials sharing the same color background and location (Fig. 4b). *Within context similarity (WCS)* was calculated with the same color background but different location, and *between context similarity (BCS)* was calculated between trials with both different color backgrounds and different locations. This was done separately for subsequently remembered and forgotten trials. Nonparametric Cluster-Based Permutation Test was used to determine the significance. Representations associated with good and poor memory states were subsequently identified by comparing the representational similarity metrics between remembered and forgotten trials. This approach enabled us to determine whether specific types of neural representations, as captured by WLS, WCS, and BCS, were predictive of successful memory encoding. By linking trial-by-trial memory outcomes with underlying neural pattern similarity, we aimed to characterize representational features which served as neural markers of memory states.

### Nonparametric Cluster-Based Permutation Test

To evaluate whether neural representation was reliably greater for remembered than forgotten trials, we conducted a nonparametric statistical test based on the cluster-level permutation^76^. For each time window, we first contrasted the two conditions using a one-sided test (remembered > forgotten). Samples exceeding a predefined threshold (*p* < 0.05) were grouped into clusters according to temporal adjacency. A cluster-level statistic was then obtained by summing the corresponding t values within each cluster.

To estimate the null distribution, condition labels (i.e., remembered vs. forgotten) were randomly reassigned across trials 1,000 times. For each permutation, the same clustering procedure was applied, and the maximum cluster-level statistic was retained to build the surrogate distribution. If no sample passed the initial threshold in a given permutation, a cluster value of zero was assigned. The observed cluster statistic was compared against this null distribution, and the empirical p-value was defined as the proportion of surrogates that yielded a cluster statistic equal to or larger than the observed one. Clusters with *p* < 0.05 were considered significant, indicating temporal regions that reliably expressed a positive SME.

### Channel-wise WLS SME analysis

To further characterize the contribution of individual channels to WLS SME, we computed channel-wise WLS SME. Unlike for the global representation approach which combined features across all channels, this analysis focused on the time-frequency features from a single channel. The time-frequency feature extraction and similarity calculation procedures for each channel followed the similar method as described above for the global representational similarity analysis (i.e., using 200-ms sliding windows with 10-ms steps, averaging spectral power per frequency within each window, and computing Spearman correlation between trials followed by Fisher Z-transform). Channel-wise similarity was calculated separately for remembered and forgotten trials. The reliability of each channel’s contribution was evaluated using a *t* test across trials.

Based on these statistics, channels were further classified into three categories reflecting the directionality of the SME: *WLS_c_ SME+ channels* (showing significantly stronger WLS during remembered than forgotten trials), *WLS_c_ SME- channels* (showing significantly weaker WLS for remembered trials), and *non-SME channels* (showing no significant difference).

### Jackknife analysis to estimate the channel contribution to WLS SME

We applied a jackknife procedure to further evaluate the contribution of each channel to the global WLS SME^77^. First, the mean WLS was calculated across remembered (*wLs_rem_all_*) and forgotten trials (i.e., *wLs_forg_all_*). Next, for remembered and forgotten trials, we computed the contribution of channel *i*. This was done by recalculating WLS after excluding that channel (i.e., *wLs_rem_all-i_* and *wLs_forg_all-i_*). Then we calculated the difference scores (i.e., *wLs_rem_all_*-*wLs_rem_all-i_* and *wLs_forg_all_*-*wLs_forg_all-i_*). These difference scores reflected how much channel *i* contributed to the global WLS in each memory state condition. Finally, we performed a *t* test comparing the distribution of difference scores from remembered trials against forgotten trials. The resulting *t* value for each channel *i* was defined as its jackknife WLS SME (Supplementary Fig. 11a). This metric quantifies whether a channel’s contribution to the global pattern was significantly different for successful versus unsuccessful encoding.

Following the jackknife analysis, channels were categorized into three groups (*WLS_c_ SME+*, *WLS_c_ SME-*, and *non-SME*), based on the same criteria and threshold as those used in the channel-wise SME analysis.

### Multi-level mediation analysis

To investigate the neural mechanisms underlying stimulation-induced memory changes, we performed a multilevel mediation analysis. The independent variables included stimulation site (between-subject factor, gray matter vs. near white matter) and stimulation frequency (within-subject factor, 0 Hz [Sham], 5 Hz, and 50 Hz). Trial-wise neural activity (theta activity/representations) served as the mediator variable and was z-scored across trials within each participant to control for inter-individual variability. The dependent variable was memory performance, coded as a binary outcome (correct vs. incorrect).

This model statistically evaluates the hypothesis that the effect of stimulation (independent variable) on memory (dependent variable) is transmitted, at least in part, through changes in neural activity (mediator variable). Following the standard mediation framework, we estimated key paths: path *a* is the effect of stimulation on the neural mediator (*β*_1_ in Eq.1), Path *b* is the effect of the mediator on memory while controlling for stimulation (*γ*_4_ in Eq.2), and Path *c’* is the direct effect of stimulation on memory after accounting for the mediator (derived from Eq.2). The total effect (path *c*) of stimulation on memory is derived from Eq.3. The indirect effect (*a*\**b*) quantifies the portion of the total effect mediated by neural activity.

The mediation analysis proceeded in three steps. First, a linear mixed-effects model (the mediator model) was fitted using *lme4* packages^78^ to estimate the effects of stimulation frequency, stimulation site, and their interactions on neural activity metrics (Eq.1).

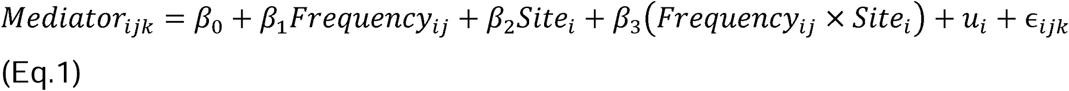

*Mediator_ijk_* was the neural activity on trial *k* of participant *i* under frequency condition *j*. *Frequency_ij_* was the stimulation frequency, whereas *site_i_* was the stimulation site category for participant *i* (gray matter or white matter).

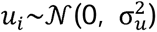 was the random intercept for participant, whereas 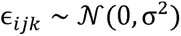 was the residual error term.

Second, a generalized linear mixed-effects model (the full model) with a logit link function was constructed to predict memory accuracy from both stimulation variables and neural activity, incorporating fixed effects for stimulation factors and the mediator (Eq.2).

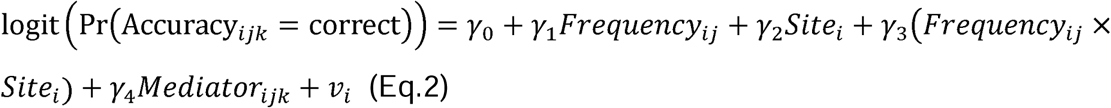

*Accuracy_ijk_* was the binary memory outcome, and 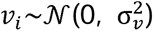 was the participant-level random intercept.

Third, to quantify the direct effect of stimulation on memory without accounting for the mediator, we fitted another generalized linear mixed-effects model excluding the mediator term (Eq.3):

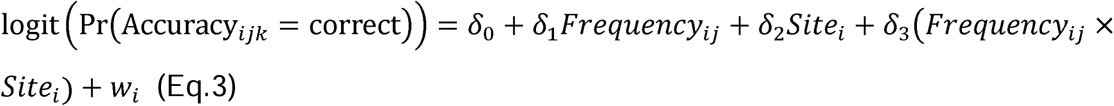

The indirect effect of stimulation on memory through neural activity was calculated as the product of the fixed effect estimates from the mediator model and the full model. To assess statistical significance, we performed non-parametric bootstrapping with 1,000 resamples. A 95% percentile-based confidence interval and two-tailed *p* values were obtained using the *boot* package^79^.

### Functional connectivity analysis

In order to obtain the connection between channels with stimulation targets, we calculated theta coherence between channel pairs during resting period^32,34^. Coherence was calculated by the cross-spectral density between signals of two channels, divided by the product of the auto-spectral power of each channel.

Continuous resting state data was firstly epoched into 2s segments with an overlap of 1s, and then we applied the multi-taper method to estimate 3-8 Hz spectral density.

Inter-channel coherence values were averaged in theta-frequency band. Given that there were anode and cathode stimulation channels, we averaged the coherence between the recording channels and the two stimulation targets.

### Bootstrap-Based Estimation of Predictor Differences

To formally compare the predictive strengths of two memory-related signatures, we employed a subject-clustered nonparametric bootstrap procedure. This analysis was designed to test whether stimulation-induced changes in theta power were more strongly predicted by the theta SME than changes in WLS were predicted by the WLS SME. Specifically, subjects were resampled with replacement to generate bootstrap datasets, while maintaining within-subject dependency structures. For each bootstrap sample, two mixed-effects models were fitted: one predicting theta power from SME and stimulation condition, and another predicting WLS and stimulation condition, each including random intercepts for subject and channel. Simple slopes were estimated using the *emtrends* function. The condition-specific slope differences between the theta and WLS models were then computed for each bootstrap replicate. Across 1,000 resamples, we derived the empirical distribution of slope differences, from which 95% confidence intervals were obtained by percentile method. Two-tailed *p* values were calculated as twice the proportion of bootstrap replicates in which the slope difference was less than zero.

## Data availability

Data and analysis scripts will be made publicly available upon publication of this manuscript.

## Acknowledgements

This work was supported by the National Natural Science Foundation of China (32330039, 32441110 and 82271494), 111 project (BP071903), and Beijing Natural Science Foundation (L256007).

